# Synthesis-driven reverse metabolomics reveals 3-hydroxy *N*-acyl amides as gut microbial molecules

**DOI:** 10.64898/2026.01.07.698252

**Authors:** Victoria Deleray, Vincent Charron-Lamoureux, Kyle Vittali, Helena Mannochio-Russo, Crystal X. Wang, Corinn Walker, Karsten Zengler, Dionicio Siegel, Pieter C. Dorrestein

## Abstract

3-hydroxy *N*-acyl amides are bioactive lipids with reported anti-obesity and glucose-regulating effects, yet they are rarely detected in untargeted metabolomics studies because they are largely absent from existing spectral reference libraries. To address this gap, we synthesized an MS2 spectral resource comprising 436 structurally diverse 3-hydroxy *N*-acyl amides, spanning 3- to 18-carbon chains with a wide range of amine headgroups such as ornithine, valine, and dopamine. Using a synthesis-driven reverse metabolomics approach, we found 161,626 spectral matches across 54,744 publicly available files in untargeted metabolomics datasets revealing widespread occurrences in biological samples, including human-derived specimens. Of these molecules detected through MS2 spectral matching, 334 represent newly reported biological entities. We further confirmed their presence in human saliva, stool, and skin using retention time and ion mobility measurements. Frequent detection in microbial datasets and validation in communities of human-derived gut bacteria support microbial production. Several metabolites also showed altered abundance in individuals with diabetes mellitus, showing that this lipid class is modulated in human metabolic disease. Together, these findings establish 3-hydroxy *N*-acyl amides as a distinct and biologically relevant lipid class, and the accompanying MS2 spectral resource will enable their broader recognition and study in untargeted metabolomics data.

## Introduction

3-hydroxy *N*-acyl amides are a class of compounds consisting of an amine head and β-hydroxylated fatty acid tail (**Fig. 1a**), and some of these structures have been described previously (**Fig. 1b**). For example, 3-hydroxyhexadecanoyl ornithine is created by *olsD* in *Burkholderia cenocepacia*^1^, a virulent pathogen affecting cystic fibrosis patients, and other ornithine β-hydroxy lipids are found in many eubacteria^2^. Two 3-hydroxy *N*-acyl amides, 3-hydroxytetradecanoyl lysine and 3-hydroxytetradecanoyl ornithine, have been detected in bovine extract as microbial-host derived products of *Neisseria meningitidis*, with stimulatory and anti-stimulatory effects on different parts of the immune system^3^. A Mosher ester analysis^4^ of those compounds revealed their S-configuration, indicating their fatty acid chain originates from β-oxidation^3^ (**Fig. 1a**), though they have never been reported in humans. 3-hydroxyhexadecanoyl ornithine is produced by *B*. *cenocepacia* and has strong potential for its use as an adjuvant for its strong immunostimulatory effects in mice^5^. Additionally, Cbeg12 encodes the production of 3-hydroxyhexadecanoyl glycine, which acts as a signaling molecule via G-protein coupled receptor (GPCR) G2A/GPR132 in *Bacteroides vulgatus* with its mammalian host^6^. β-hydroxybutyryl-leucine, -methionine, -phenylalanine, and -valine have been identified as products of ketogenesis in mice via the CNDP2 enzyme^7^ and found in humans, an important gap given the known relationship between microbial communities and host energy metabolism^8^, and the known bacterial origin of this chemical moiety. The structural and functional characteristics of these 3-hydroxy *N*-acyl amides are vast and remain unexplored in the context of human metabolism and microbiome intervention. A summary of the known 3-hydroxy *N*-acyl amides can be found in **Fig. 1b**.

**Fig. 1.**
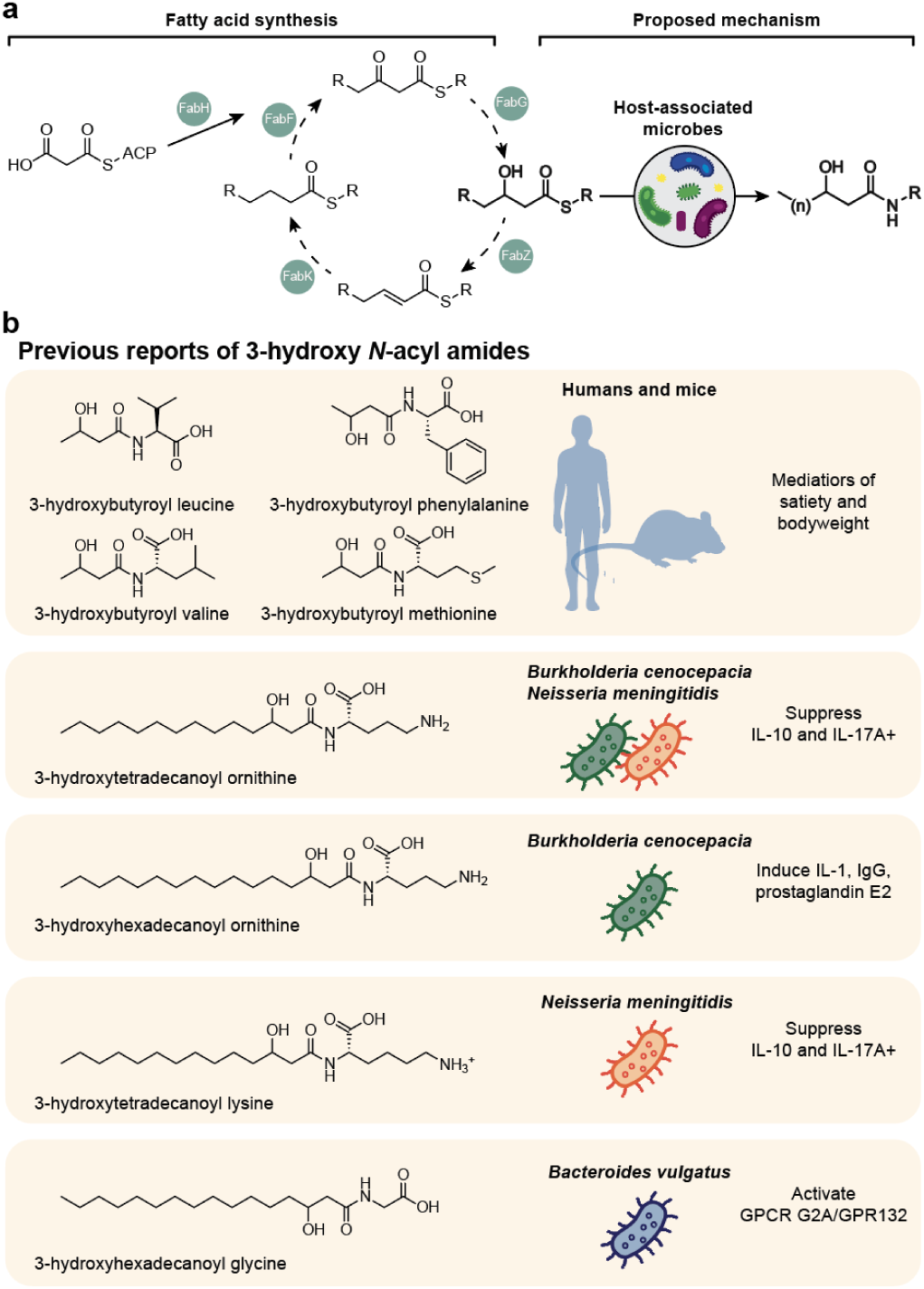
Production and functions of 3-hydroxy *N*-acyl amides. **a.** Summary of known metabolic pathway of the synthesis of fatty acids to 3-hydroxy-fatty acyls occurring in the mitochondria^11^. Our proposed mechanism shows the 3-hydroxy fatty acyl undergoing alternative metabolism to become a 3-hydroxy *N*-acyl amide. **b.** Summary of previous reports of 3-hydroxy *N*-acyl amides, including structure, name, and organism it is found in, and known activities of these molecules (*in vivo* or *in vitro*) are referenced in the introduction.

Structures or tandem mass spectra (MS2) of 3-hydroxy *N-*acyl amides are scarce in reference spectral libraries. Despite their important functional roles, these metabolites are largely overlooked in untargeted metabolomics experiments. For example, collated resources of MS2 libraries such as MassBank of North America (MONA)^9^ and Global Natural Product Social Molecular Networking (GNPS)^10^ contain no 3-hydroxylated *N*-acyl amides.

Other *N*-acyl lipids have recently been unveiled as key microbial signaling molecules through a reverse metabolomics strategy^12,13^. Compared to traditional mass spectrometry-based metabolomics, where experiments are designed prospectively, samples are acquired, MS/MS spectra are generated, and the data are searched against known libraries to assign a structure, reverse metabolomics is a method that searches an MS2 spectrum across entire untargeted metabolomics repositories such as GNPS/MassIVE, Metabolights, Metabolomics Workbench, and NORMAN^14^. Using data science and integrating metadata information, any MS/MS spectra can be linked to biological-rich information such as organ/biofluids distribution, health phenotype association, as well as sex and age^15^. This was done previously to investigate organ distribution and extraction solvent in a collection of (non-hydroxylated) *N*-acyl amide spectra that were obtained from either combinatorial synthesis or public data mining using mass spec query language (MassQL); then, their biology was characterized within thousands of public untargeted metabolomics datasets^13^.

Our approach leverages the synthesis-based approach^12,16^, which creates both a spectral and physical library of these 3-hydroxy *N*-acyl amides. Using the MS2 spectra of these novel synthetic compounds, we mapped their occurrence in humans and microbial monocultures, and confirmed their presence in humans with orthogonal validation methods, and microbial origins using *in vitro* culturing of gut-derived bacteria. With this approach, we have newly annotated thousands of MS2 spectra within human samples, bacteria, and other organisms, and further confirmed (Metabolomics standards initiative level 1^17^) nine previously unreported 3-hydroxy *N*-acyl amides within various human tissue/biofluids and microbial cultures. Curation of this library led to an increased understanding of energy metabolism and microbial transformation within human microbiomes, and serves as a valuable resource for future mechanistic studies.

## Results and Discussion

### Curation of the spectral library and pan-repository analysis

At present, the average annotation rate of all MS2 using publicly accessible reference libraries in the major repositories Metabolights, Metabolomics Workbench, and GNPS/MassIVE is roughly 14%^16^. It is likely that many of the unannotated MS2 spectra belong to 3-hydroxy *N*-acyl amides, but have gone unannotated due to a lack of reference libraries. To facilitate their annotation, we aimed to create a reference library by performing 55 multiplexed reactions containing mixtures of 19 3-hydroxy fatty acids, and 42 amine head groups **(Supplementary Table 1)**. Each multiplexed reaction had one to three 3-hydroxy lipids and was reacted with 4 to

24 different amines to generate their corresponding amides. The head groups included proteinogenic amino acids (e.g., valine, tryptophan, alanine), polyamines (e.g., putrescine, cadaverine, ornithine), and amino acid metabolites (e.g., citrulline, serotonin, and dopamine). The lipid tails were chosen based on commercial availability and reasonable costs, and amines were chosen based on availability in the Dorrestein/Siegel laboratories. Theoretically, these reactions could yield 798 distinct 3-hydroxy *N*-acyl amides, of which 536 were detected by liquid chromatography tandem mass spectrometry (LC-MS/MS).

To add confidence in the generated library, we filtered the data through SIRIUS^18^, a software which utilizes isotope patterns and fragmentation trees to aid in substructure annotation, to ensure the MS2 data could be explained by the assigned structures. Manual inspection of diagnostic fragment peaks generated an MS2 library of 436 unique compounds represented by 726 MS2 spectra. The distribution of adducts were [M+H]^+^ (n=416), the [M+Na]^+^ (n=147), [M+K]^+^ (n=25), [M+NH4]^+^ (n=53) adducts and MS2 of water loss in-source fragment [M-H_2_O+H]^+^ (n=122) were added to the public MS2 libraries in GNPS and in Zenodo^19^ as “3-HYDROXY-ACYL-AMIDES-LIBRARY”. We manually curated the MS2 spectra by looking for the diagnostic peaks of the head group. For example, to validate the spectra for glutamic acid conjugated to a C12:0 chain (12 carbons and 0 double bonds), we used data from MS2 spectra on the [GNPS library](https://gnps.ucsd.edu/ProteoSAFe/gnpslibrary.jsp?library=GNPS-LIBRARY) where two public spectra for glutamic acid show consensus fragment peaks of 84.04, 102.06, and 130.05. Since these three peaks were also found in the spectra assigned to the 3-hydroxy-conjugate (3-hydroxylauroyl glutamic acid), they were retained during filtering. Most of these *N*-acyl amides for which we created a reference library did not only have MS2 spectra in the public domain but were also largely absent from the most comprehensive structural databases such as LipidMAPS^20^ and PubChem^21^.

Only three of these *N*-acyl amides were found in the lipid database, LipidMAPS^20^, the leading structural resource in the lipid field. These were 3-hydroxyoctadecanoyl ornithine, 3-hydroxyhexadecanoyl ornithine, and 3-hydroxytetradecanoyl glycine. Only 79 were found in PubChem^21^, and only one, 3-hydroxyoctadecanoyl glycine has previously been reported to be detected by the human metabolome database (HMDB)^22^, and none in the MassBank of North America (MONA)^9^ (**Supplementary Table 2**). This new spectral resource represents a unique opportunity to reveal detected but not yet annotated biochemistry in the public domain data. We also aim to improve annotation in future LC-MS/MS-based untargeted metabolomics studies, leading to an enhanced understanding of biology.

To provide evidence of their presence in biology, the reference library consisting of 763 M/MS spectra underwent reverse metabolomics analysis (**Fig. 2a**). Using reverse metabolomics, we found 161,626 spectral matches in the public domain with a minimum cosine similarity of 0.7 using a minimum 3 matched ions, representing 334 unique 3-hydroxy *N*-acyl lipids (**Fig. 2b**). These were matched to data from 1,255 studies (GNPS/MassIVE n=1,192, Metabolights n=45, Metabolomics Workbench n=19) in 54,744 data files. Of the unique data files, 1,025 match to Metabolights, 59 to Metabolomics Workbench, and 53,660 GNPS/MassIVE **(Supplementary Table 3)**.

**Fig. 2.**
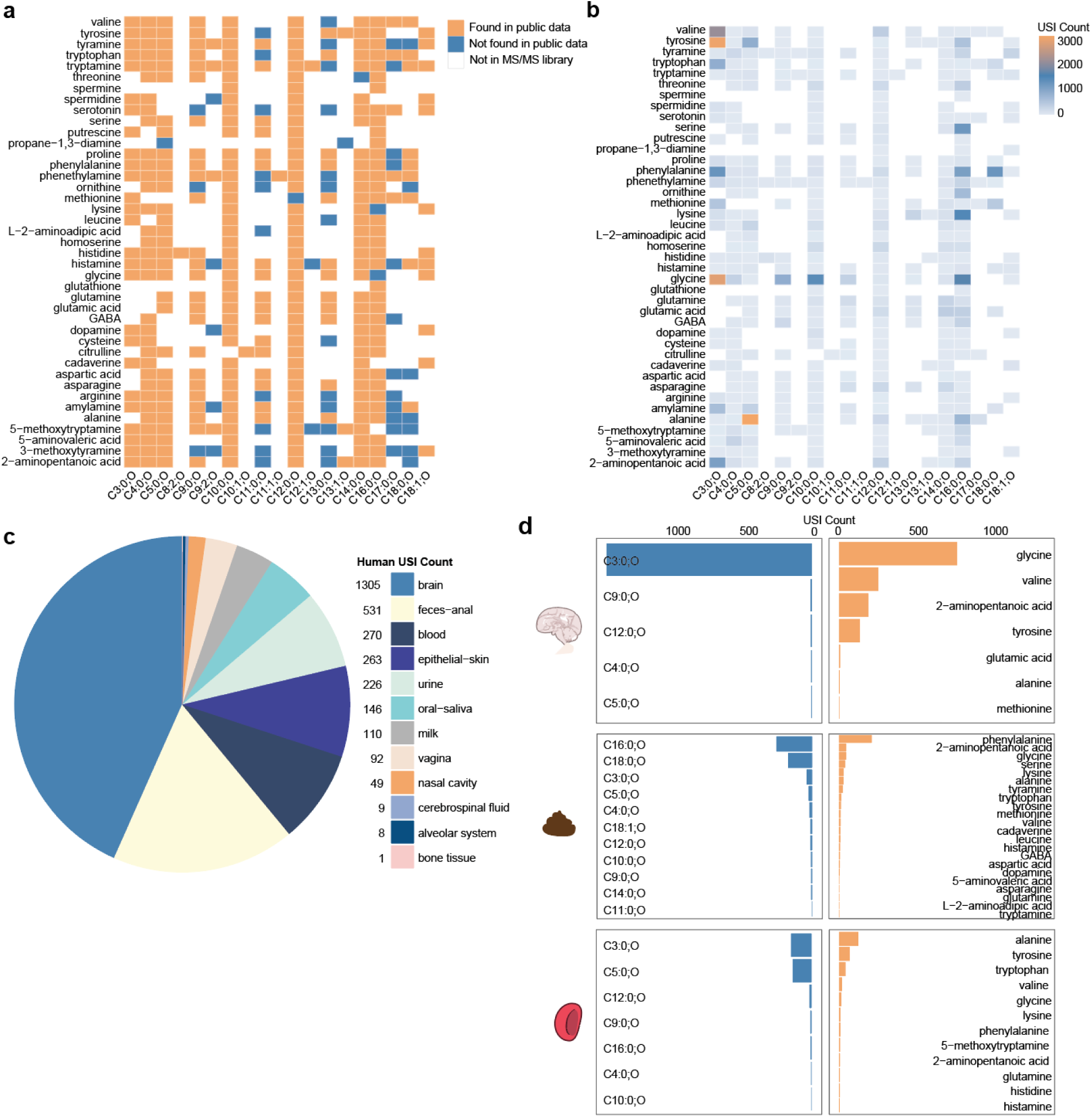
Pan-repository analysis of the 3-hydroxy *N*-acyl amide library. **a.** Heatmap showing presence/absence of 3-hydroxy acyl amide combinations of differing tails (chain lengths) and heads (amines). Lipid chains are annotated as the number of carbons and number of unsaturations in the chain, followed by annotation of a hydroxyl modification. The y-axis shows head groups used in synthesis ordered alphabetically, and the x-axis shows chain lengths used in synthesis ordered by length. A colored square indicates that a library spectrum exists for a unique head-tail combination. A blue square indicates no spectral match (minimum cosine of 0.7) exists within public data repositories for that acyl amide combination. An orange square indicates at least one spectral match (minimum cosine of 0.7) within public data. A white square indicates that the head-tail combination is not present in the MS/MS library. **b.** Heatmap represents the number of MASST spectral matches (minimum cosine of 0.7) to each library spectrum subset by head and tail. **c.** Pie chart shows the distribution of the entire 3-hydroxy acyl amide library against all public data, with metadata labeled as “Homo sapiens” with sample type (UBERONBodyPartName) information found in Pan-ReDU^13^. The sample type with the most spectral matches is brain, followed by feces-anal (concatenation of samples labeled feces or anal tissue). **d.** Bar charts show the number of spectral matches per tail and head substructure for the top 3 most annotated sample types-brain, feces-anal, and blood (includes serum and plasma).

The proposed origin of these lipids is β-oxidation or classical acetate-mediated lipid assembly, where fatty acids are built up or broken down in two-carbon units^11,23^. When comparing the unsaturated lipids to their saturated counterparts for which we had spectra available (e.g., 3OH-C10:0 to 3OH-C10:1), there was a decrease in numbers (matching unsaturated n=3,933 and saturated n=89,094). However, even chains (n=45,531) showed similar prevalence to odd chains (n=43,563) by spectral matching. A significant portion of the odd chains found were C3:0 (n=17,893, representing 41% of the matches). It is likely that these conjugates’ lipids originate elsewhere, or the matches come from an isomer such as lactate.

Among the samples in Pan-ReDU^14^, the infrastructure which harmonizes metadata from four main mass spectral repositories into controlled ontologies, we obtained spectral matches to 3,010 human data files, 2,386 to rodents, 4,278 to plants^24^, 6,505 to microbes^25^, 615 to food^26^ **(Supplementary Table 4)**. The remaining data files belonged to other organisms or did not yet have computer-readable metadata. When comparing the number of unique compound matches to each domain, microbes had the largest number of matches to MS2 of the 3-hydroxy *N*-acyl amide library, indicating that many of these molecules were likely of microbial origin.

Of the available datasets in rodents, the whole family of compounds was found in the blood (1,137 MS2 matches, representing 9 compounds), feces (401, 19 compounds), caecum (316 MS2 spectra, 8 compounds), ileum (144 MS2 spectra, 10 compounds), and jejunum (130 MS2 spectra, 4 compounds), among other tissue types (**Supplementary Table 5**). In humans, MS2 detection was in brain (1,305 MS2 spectra, representing 9 compounds), feces and anal (531 MS2 spectra, 42 compounds), blood (270 MS2 spectra, 16 compounds), skin (263 MS2 spectra, 14 compounds), and urine (146 MS2 spectra, 12 compounds) (**Fig. 2d**; **Supplementary Table 6**). In humans, the fecal samples had the highest diversity of 3-hydroxy *N*-acyl amides containing combinations of 11 3-hydroxy fatty acids and 22 amine head groups, whereas the brain had the most MS2 matches with a unique combination of only 5 tails and 7 heads through the overall repository-scale searches. It is important to note that the repository-scale analysis is based on spectral matching alone, and therefore, other potential isomers might exist. To reach level 1 confidence according to the metabolomics standards initiative^17^, we used retention time and drift time comparisons in addition to spectral matching in select datasets.

### Bacterial origin of 3-hydroxy *N*-acyl amides

Searching our 3-hydroxy *N*-acyl amide library against public data revealed that most matches were against microbial samples, with many found in *Burkholderia*, *Bacteroides*, and other eubacteria^1,2,6^ by using a domain-specific MASST, microbeMASST, with 61,000 taxonomically defined microbial data of monocultures^25^. We found 6,505 MS2 matches from the library to the microbial data, representing 135 3-hydroxy *N-*acyl amide spectra belonging to 111 unique compounds (**Fig. 3a**).

**Fig. 3.**
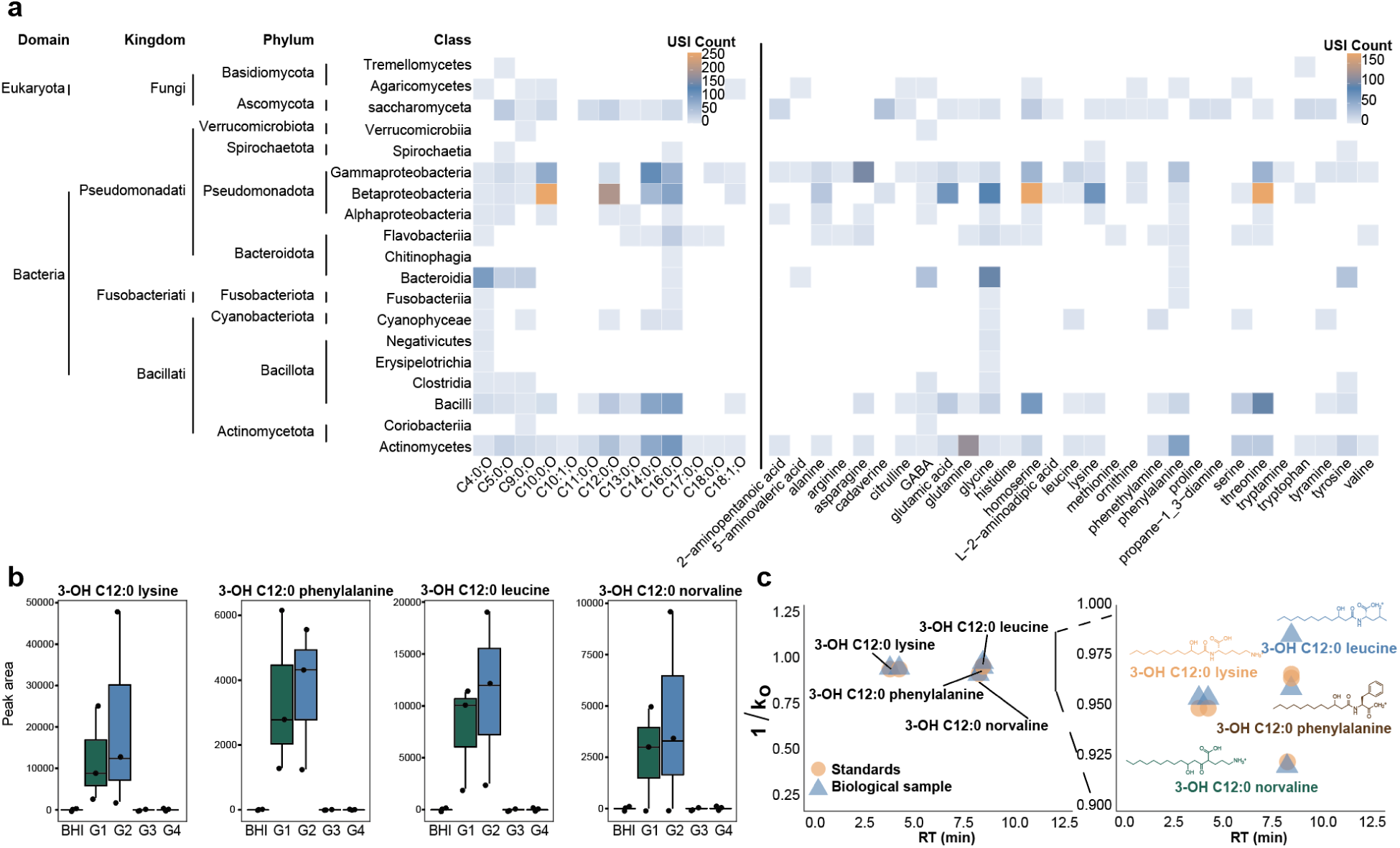
Bacterial origin of 3-hydroxy *N*-acyl amides. **a.** Heatmap shows the number of USI (public MS2 spectra) which had a microbeMASST^16^ spectral match to the 3-hydroxy *N*-acyl amide library stratified by the head (amine) and tail (chain length) of the molecule. The x-axis is clustered based on chain length from smallest to largest (left panel) and amines by alphabetical (right panel). The y-axis is clustered by NCBI taxonomy and labeled from domain to class. **b.** Boxplots show feature abundance of detected 3-hydroxy *N*-acyl amides within the synthetic microbial communities experiment at T=72 h. BHI represents the media with the addition of 3-hydroxy lipids without any added bacteria. G1 represents fast growing gut community members, G2 represents fast-medium growing, G3 represents medium-slow, and G4 represents slow growers. Fast growers and medium-fast growers show an increased abundance of the acyl amides compared to the media control. MS2 mirror plots with the library can be found in **Supplementary Table 8**. **c.** Summary of retention time and drift time (1/K_0_) peaks of detected 3-hydroxy *N*-acyl amides within the microbial culturing experiment.

Many of the bacteria within microbeMASST, which had matches to the 3-hydroxy *N*-acyl amide library, belonged to classes of bacteria that are part of the human gut microbiome (e.g., Bacilli, Clostridia)^27,28^. We therefore hypothesized that many of the 3-hydroxy *N*-acyl amides detected in human or mouse (**Supplementary Table 5, Supplementary Table 6)** might be a product of microbial transformation of the proposed starting materials-3-hydroxy lipid chains, which are a product of endogenous lipid metabolism **(Fig. 1a)**, and various head groups, including amino acids and other endogenous small amines. To test this hypothesis, we cultured defined gut communities^29^ in brain heart infusion (BHI) media, which contains a broad range of substrates including amino acids^30^, and we added seven 3-hydroxy lipids, including C3:0, C4:0, C5:0, C11:0, C12:0, C13:0, and C16:0 based on their availability in our lab **(Supplementary Table 7)**. At 72 h, we observed that fast-growers (G1) and medium-fast grower (G2) communities were able to produce some 3-hydroxy *N*-acyl amides, most notably C12:0 chains, that were not observed in the control (medium alone) **(Fig. 3b**). We confirmed their identity through spectral matching, retention time, and drift time analysis **(Fig. 3c, Fig. S2a)**, confirming the production of these molecules by bacteria.

### Potential origin of 3-hydroxy *N*-acyl amides in humans and their verification

We assessed the presence of these 3-hydroxy *N*-acyl amides within various human datasets found in Pan-ReDU, in different biological conditions utilizing the MS2 spectral library. We first investigated distribution in disease states, an approach to begin to understand the origin or function of these metabolites. There were 159 spectral matches to samples labeled as “inflammatory bowel disease” (IBD), 120 spectral matches to “primary bacterial infectious disease” (PBID), 60 to “diabetes mellitus” (diabetes), and others (**Supplementary Table 9**). Investigation of the IBD and PBID datasets showed no metadata available to compare the abundance of the 3-hydroxy *N*-acyl amides between healthy and diseased participants. We thus targeted diabetes, whose datasets had metadata for urine samples from diabetics and non-diabetics. We performed feature extraction and feature-based molecular networking to the raw data available in MassIVE (MSV000082261). We found a spectral match to the library belonging to 3-hydroxylauroyl glutamic acid, then propagated annotations within the molecular network using spectral similarity, mass defects, and retention time, which putatively identified other hydroxy-lipid glutamic acid conjugates **(Fig. 4a)**. Analysis of the peak areas of these features showed significant (*p* < 0.05) depletion of these metabolites in the urine of diabetic patients **(Fig. 4b)**. Metabolic dysfunction, which may impact the synthesis or β-oxidation of fatty acids, is a characteristic of type 1 diabetes^31^. If indeed the lipid substrates of 3-hydroxy *N-*acyl amides originate from this endogenous pathway, we would expect to see dysregulation of *N-*acyl amides in diabetics as we do here. Alternatively, some connections between microbiome composition and type 1 diabetes have been found, which may also contribute to this pattern^32–34^.

**Fig. 4.**
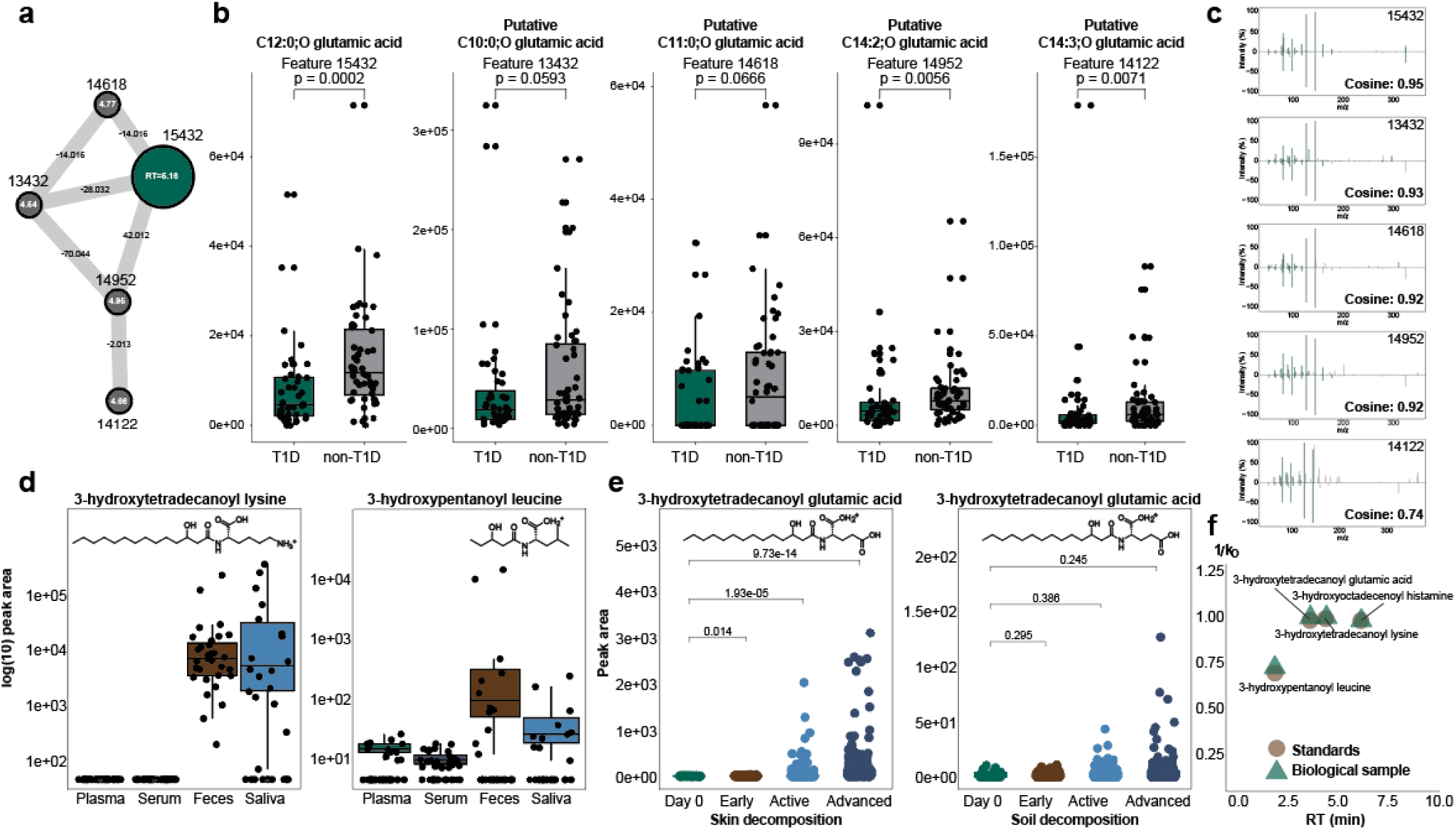
The production of 3-hydroxy *N*-acyl amides depends on energy metabolism, organ distribution, and microbial activity. **a.** Network of features in the feature-based molecular networking shows related spectra of 3-hydroxylauroyl glutamic acid and their relative delta masses and retention times. **b**. Univariate analysis of selected feature abundances from dataset MSV000082261. Feature 15432 was annotated through MS2 spectral matching (metabolomics initiate level 2) as 3-hydroxylauroyl glutamic acid, which is significantly (p<0.05) depleted in urine samples labeled type 1 diabetes mellitus in the public metadata compared to the healthy samples. Features related to 3-hydroxylauroyl glutamic acid (13432, 14618, 14952, 14122) are significantly (p<0.05) depleted in the type 1 diabetes mellitus samples. Significance was assessed using the Wilcoxon rank-sum test. **c.** MS2 spectral mirror plots show high cosine similarity (Cosine = 0.95) between feature 15432 and the synthetic spectra of 3-hydroxylauroyl glutamic acid (CCMSLIB00013641087). Other selected features in the molecular network (panel b) have high cosine similarity (>0.7) to feature 15432, showing their structural similarity. **d.** Boxplots show feature abundance (log10 fold) of detected 3-hydroxy *N*-acyl amides, 3-hydroxytetradecanoyl lysine and 3-hydroxypentanoyl leucine, within different sample distributions in the MSV000094097 dataset. **e.** Boxplots show feature abundance of 3-hydroxytetradecanoyl glutamic acid within the MSV000084322 dataset sampling skin swabs and soil swabs of outdoor body decomposition over time. 3-hydroxytetradecanoyl glutamic acid was significantly increasing in skin decomposition (p<0.05) with time. No significant increase in soil, because the feature abundance was low (<150 peak area). Significance was assessed using the Wilcoxon rank-sum test. The boxplots represent first (lower), interquartile range (IQR), and third (upper) quartile. **f.** Summary of retention time and drift time (1/K_0_) peaks of detected 3-hydroxy *N*-acyl amides within all public data experiments. MS2 mirror plots with the library can be found in **Supplementary Table 8.**

Re-analysis of data samples from a cohort of osteoarthritis patients showed some tissue specificity of two newly annotated 3-hydroxy *N*-acyl amides **(Fig. 4d)**. 3-hydroxypentanoyl leucine was present in feces, serum, plasma, and saliva samples in this cohort. 3-hydroxytetradecanoyl lysine, however, was present in many samples across only fecal and saliva samples. These molecules were also validated with retention time and drift time analysis (**Fig. 4f**, **Fig. S1b**). This tissue/biofluid specificity supports the conclusion that these molecules are indeed microbially made, as the gut and salivary microbiome are well documented^35^. Reanalysis of a dataset which sampled human cadavers’ skin and surrounding soil composition showed the abundance of 3-hydroxytetradecanoyl glutamic acid significantly increased (*p* < 0.05) with advancement of skin decomposition (**Fig. 4e**). This molecule was present in very low or no abundance (peak area <150) in the soil and had no significant (*p* < 0.05) changes with decomposition time. This molecule was validated with retention time and drift time analysis (**Fig. 4f**). 3-hydroxytetradecanoyl glutamic acid increases with decomposition, mirroring the increase in microorganisms and their products^36^, serving as further evidence for microbial origin of this molecule. Another undescribed molecule, 3-hydroxyoctadecenoyl histamine, was detected and validated in additional fecal sample part of a cohort from the HIV Neurobehavioral Research Center (HNRC)^13^, which aims to investigate neurological complications in people with HIV **(Fig S1b)**.

### Conclusion and future outlook

This high-throughput generation of MS2 spectra of 3-hydroxy *N*-acyl amides resulted in an expansion of untargeted metabolomics annotation and functional and spatial hypotheses generated for these underreported compounds. By targeting a class of compounds with known functional characteristics such as amide linkages and β-hydroxylations, we were able to increase the known diversity of human-associated metabolites. This study highlights the importance of reanalysis of public data, a vastly growing resource of thousands of study designs, models, and instrumentation. We show that this reverse metabolomics method can contextualize the source or function of individual molecules by utilizing available data. We found that this class of 3-hydroxy *N*-acyl amides is found in hundreds of LC-MS/MS studies among diverse types of organisms and human samples. We demonstrated that community cultures of gut bacteria, when given 3-hydroxy lipids as substrates, can lead to the production of *N*-acyl amides. Metabolic dysfunction influences the abundance of some 3-hydroxy *N*-acyl amides, and we provided insight into host-microbe metabolic interactions, though pathways are out of the scope of this study. Fundamentally, to propose such interactions and functions, we must be able to detect and annotate these compounds, which our public library will grant.

We shared 765 unique MS2 spectra in this library, but thousands of other amine heads and unique lipid chain lengths remain uninvestigated. Further synthesis of such compounds, or other *in-silico* or computational techniques are needed to continue the investigation of such compounds. We hope that the workflow used here and the knowledge of the fragmentation patterns of these compounds can be leveraged to expand this understanding and increase general metabolomics annotations in a wide variety of experiments globally.

We hope this resource becomes widely used and incorporated into untargeted metabolomics workflows and is made simple by its addition in the GNPS library and Zenodo. However, the discovery and characterization of these molecules through this resource has limitations.

Many ion forms, or in-source fragments such as water loss, of one molecule exist in mass spectrometry data. This library captured [M+H]^+^ for all library compounds and other common ions such as [M+NH4]^+^, [M+Na]^+^, [M+H-H2O]^+^, or [M+K]^+^. This library cannot comprehensively annotate all ion forms of one molecule, some of which are biased to different analytical methods or instrumentation. Some computational methods have been developed to aid researchers with detection of all ion forms^37^.

Similarly, with all the different instrumentation and analytical methods that exist for LC-MS/MS workflows^14^, it is likely that the pan-repository search was not comprehensive of all 3-hydroxy *N*-acyl amides in the library, and the results are likely biased to similar methods used in the creation of the library. As such, results are likely to change with the increase of data deposition in the repositories used here. We recognize that obtaining spectra from multiple instruments or analytical methods, such as negative ionization mode, would improve the library and serve as a future direction.

For the spectral matches we did obtain, interpretation of results is limited to the metadata which is publicly available or documented. We aim to use the metadata to give insight into the potential origin of the molecule by recognizing general patterns, such as several pieces of evidence of microbial production.

Lastly, we utilized a multiplex synthesis method for the high-throughput curation of the MS2 spectral library. The extraction of the spectra and annotation of the 3-hydroxy *N*-acyl amides of the reaction mixtures was aided by two annotation tools and manual inspection of the spectra for the correct structural annotation. We did not produce isolated and purified compounds to validate the spectra, nor obtain NMR or other information to validate the synthesis.

## Methods

### Multiplexed Reactions

Separate 4 reagent substrate mixtures were created for the multiplex synthesis reactions (**Supplementary Table 10**). In 20mL of LC-MS grade water, we added 8 umol of each amine and stored at –80°C until the reactions were performed. Either 1, 2 or 3 3-hydroxy fatty acid(s) (1eq.) and 2mL of THF were added to a 20mL scintillation vial with a magnetic stir bar. To this solution, neat ethyl chloroformate (1.2 eq.) and triethylamine (1.2 eq.) were subsequently added, and the reaction flask was placed in an ice water bath. In 1.5h, 1mL of amine mix was added, and the reaction continued to stir with ice cooling, allowing the ice to melt over 12h. LC-MS/MS analyses were achieved as follows using 1 µL of the reaction mixture diluted in 1 mL of LC-MS grade MeOH.

### Curation of the 3-hydroxy *N*-acyl amide library

The MS2 library generation of the 3-hydroxy *N*-acyl amides were performed on a Thermo Scientific Orbitrap Exploris 240 mass spectrometer with a Thermo Vanquish UHPLC system. Samples were ionized by electrospray ionization (ESI) in positive mode using a H-ESI source (spray voltage: +3500 V; sheath gas: 50; auxiliary gas: 10; sweep gas: 1; ion transfer tube temperature: 325 °C; vaporizer temperature: 350 °C). Chromatographic separation was carried out on a Phenomenex Polar C18 column (2.6 µm particle size, 100 × 2.1 mm) using a 12-min linear gradient at 0.5 mL/min with mobile phases consisting of H2O 0.1% formic acid (A) and ACN 0.1% formic acid (B). The gradient is 0-0.5 min, 5% B; 0.5-7.5 min, 5-40% B; 7.5-9 min, 40-99% B; 9-10 min, 99%B; 10.1-12 min, 5%B. Full scans were acquired from *m/z* 80-1000 at 120,000 resolution, automatic gain control (AGC) target set to Standard. Data-dependent acquisition (DDA) selected top 10 precursors per cycle using an isolation window of *m/z* 1 and stepped HCD collision energies of 25, 35, and 50. MS/MS spectra were acquired at 60,000 resolving power with an AGC target set to Standard and automatic maximum injection time. A 3 s exclusion duration and 3 ppm mass tolerance for dynamic exclusion. Isotope exclusion was enabled. Raw data was converted to open-source “.mzML” format in PreoteoWizard^38^ MSConvert program with centroid peak picking.

The spectral library was created in the reverse_metabolomics_create_library_workflow^16^ in GNPS2 which searches for a given molecular formula in the mzML files. The molecular formulas were given as SMILES of the targeted compounds. Adducts of [M+H]^+^ were required for each compound and additional adducts [M+Na]^+^, [M+NH4]^+^, [M-H2O+H]^+^, [M+K]^+^ were included in the library if [M+H]^+^ or [M+NH4]^+^ was also included. Multiple adducts were included to enhance the rate of spectral matches to untargeted metabolomics data. The target list included 967 compounds, including the 798 acyl amides with some isomer forms when the amine head included more than one primary amine functional group. The formula search looked for ions within 10 ppm of the given formula mass given that the intensity of the MS1 ion is minimum 5e4. Further, the feature, or MS2 must have a minimum feature height of 1e5. This search found 536 unique target compounds and 483 of the compounds had an [M+H]^+^ ion form. It further filters these matches with an “MS2 explanation score” which searches all fragments of the MS2 for a formula-based substructure of the given parent formula with a minimum cosine level of 0.6. This filtering selected 443 unique compounds to create a library of 925 spectra.

Then, the spectra collected from the GNPS2 workflow are further validated for sub-structure annotation in SIRIUS 6.0.7^18^. Using the molecular formula prediction, CSI:finger ID and database search of the spectral files using the same SMILES-based target list, SIRIUS was able to find 88.2% of the compounds from the GNPS2 workflow (**Supplementary Table 11**). It found 25 unique compounds that GNPS did not. Any compound found in both GNPS2 and SIRIUS was uploaded to the GNPS public library and Zenodo^19^. Any feature found in only one of the programs was inspected manually. Acyl amides have a predictable fragmentation pattern in positive mode which include at least two high intensity peaks from the head group. Using fragmentation patterns about the head group compounds in GNPS libraries, 32 of the 52 compounds not found by SIRIUS were recovered and 11 of the 25 compounds not found by GNPS were recovered. A total of 313 compounds could not be detected in either GNPS2 or SIRIUS because of unsuccessful synthesis. This left 944 total spectra which were then manually filtered for quality to create a spectral library of 764 spectra of 436 unique compounds with unique USI values. The filtered library spectra were uploaded to the public GNPS library through the batch annotator workflow on GNPS and is named 3-HYDROXY-ACYL-AMIDES-LIBRARY.

### Pan-repository compound search

We conducted a pan-repository-scale search using a fast MASST^39^ (FASST) search, an updated and faster version of the Mass Spectrometry Search Tool (MASST), against all the public domain data that were indexed in GNPS/MassIVE, Metabolomics Workbench, and Metabolights. This search is based on the cosine similarity of the queried spectra against the ones from the public domain in GNPS/MassIVE, regardless of the instrument used for data acquisition. This search was conducted in the GNPS2 workflow fasst_batch_workflow which took the USI^40^ of the library compounds as an input, requiring matches to public data to have a raw cosine similarity score >0.7, and precursor ion and fragment ion tolerances within 0.02 Da as done in previous pan-repository analyses. Upon the initial MASST query of the entire spectral library to all three public data repositories (*metabolomics_pan_repo_index_nightly*) there are 3,358,736 spectral matches. Then, inspection of the cosine similarity score of the raw matches kept 269,049 spectra with raw cosine > 0.7 taken from the JSON of each spectral match on the metabolomics resolver^41^ on GNPS.

Domain-specific MASSTs were conducted with the REST web API using *metabolomics_pan_repo_index_nightly*^14^. One search curated 4 domain-specific output tables that contain spectral matches to specific data files in the public domain that can be mapped to the curated list of taxonomy/ontology-defined metadata. (1) microbeMASST^25^: merges the MASST spectral matches against a curated database of more than 60,000 LC-MS/MS files of microbial monocultures that were taxonomically defined; (2) plantMASST^24^: merges the MASST matches against 19,075 LC-MS/MS files of plant extracts of taxonomically defined plants and (3) foodMASST^26^: merges MASST matches against ∼3,500 LC-MS/MS files of foods and beverages categorized within a food ontology, collected as part of the Global FoodOmics project. We derived phylogeny from the NCBI taxonomy in microbiomeMASST by querying the NCBI API using R package Taxize (version 0.10.0). Creation of the tree was done through manual query of the NCBI taxonomy database.

To explore the molecules organ, tissue, and biofluid distribution in human and mice studies, we leveraged the Pan-ReDU curated metadata to filter for 9606|Homo sapiens or rodent-related datasets (“10088|Mus”, “10090|Mus musculus”, “10105|Mus minutoides”, “10114|Rattus”, “10116|Rattus norvegicus”) as written in the Reverse Metabolomics protocol^15^. Packages used for data table manipulation included readr (version 2.1.5), stringr (version 1.5.2), dplyr (version 1.1.4), tibble (version 3.3.0), tidyr (version 1.3.1), and data.table (version 1.17.8). Heatmaps, bar plots, and pie charts of the pan-repository data were created in RStudio with the ggplot package (version 4.0.0).

### Reanalysis of Public Data

The 3OH-acyl amide library was used to analyse datasets MSV000082261 (diabetes), MSV000094097^42^ (osteoarthiritis), MSV000092833^13^ (HNRC), and MSV000084322^36^ (dead bodies). For each dataset, the files were downloaded from GNPS/MassIVE and processed in MZmine^37^. The linked parameters and software version used for each study is available in **Supplementary Table 12**. The corresponding “iimn” (ion identify molecular networking) files were used in the feature-based molecular networking available on GNPS2 with library search against 3-HYDROXY-ACYL-AMIDES-LIBRARY. The networking jobs are available in **Supplementary Table 13**. The parameters for the match were set as 0.02 Da and cosine similarity score >0.7. The number of matching peaks was set to 2, as some of the spectra, like 3-hydroxybutyroyl glycine, or [M+Na]^+^ adducted ions have very few fragments in the MS2 spectra. Manual inspection of the matched spectra was used to verify the library matches to the biological spectra, and further validation through retention time and drift time matching was used if we had access to the samples. Molecular networking on the GNPS2 interface was used to find chain length analogs to the compounds detected with the library search, as used in the analysis of MSV000082261 and were labeled as putative because they are an MSI level 2. The resulting feature tables were used for relative peak area analysis. Packages used for data table manipulation included readr (version 2.1.5), stringr (version 1.5.2), dplyr (version 1.1.4), tibble (version 3.3.0), tidyr (version 1.3.1), and data.table (version 1.17.8). Box plots and heatmaps were created using the ggplot package (version 4.0.0), ggsignif (version 0.6.4), and scales (version 1.4.0) in RStudio. Statistical significance was assessed using the Wilcoxon rank-sum test. All boxplots represent first (lower), interquartile range (IQR), and third (upper) quartile.

### Bacterial cultures screening

A total of 80 strains of bacteria (**Supplementary Table 14**) were inoculated from glycerol stocks and cultured anaerobically in filter-sterilized modified brain-heart infusion (BHI) broth (75% N_2_, 20% CO_2_, and 5% H_2_)(**Supplementary Table 15**). Each strain was normalized to OD_600_ = 0.02 and combined into a master mix according to growth rate classifications: Group 1 (fast-growers), Group 2 (medium-fast growers), Group 3 (medium-slow growers), and Group 4 (slow-growers). The four groups were pooled together and diluted 1:10 to construct the full hcom (human gut synthetic community). The 3-hydroxy lipids were added at 100 µM to a shallow 96 well-plate followed by the microbial communities to be incubated anaerobically at 37 °C for 72 h. After incubation, bacterial cells were extracted with a ratio sample to solvent of 1:4 with either 80% of pre-chilled MeOH/H_2_O with 1 µM sulfamethizole and incubated overnight at 4 ℃. Samples were centrifuged at 1800 rpm for 10 min and the supernatants were transferred to a new plate before being dry in a CentriVap overnight. Samples were stored at -80 ℃ until LC-MS/MS analysis. Samples were resuspended in 50:50 MeOH/Water with 1 µM sulfadimethoxine internal standard and run on an Orbitrap Exploris 240 instrument using electrospray ionization (ESI) with chromatographic separation through the ThermoFisher Vanquish attached to a Phenomenex 2.6 µm Polar C18 100 x 2.1mm column. The synthetic samples were eluted on a 12 min gradient with flow rate of 0.5 µ/min and the mobile phase included H2O + 0.1% formic acid and ACN + 0.1% formic acid. Raw data was converted to open-source mzML format in ProteoWizard^38^ MSConvert program. The resulting .mzML files were processed in MZmine4^37^ (hcomm_batch.mzbatch at https://github.com/victoriadeleray/3OH_N-acyl_amides). The resulting feature table and spectral list (available on https://github.com/victoriadeleray/3OH_N-acyl_amides as hcomm iimn_gnps.csv and hcomm iimn.mgf) were used for relative peak area analysis and annotation within the feature-based molecular networking workflow on GNPS2 (task id=abcfa3d422534d86b2e71a24efbd7fc6). The peak area table was filtered to remove blanks, and internal standards were used for quality control. Box plots and heatmaps were created using the ggplot package (version 4.0.0) and tidyverse package (version 2.0.0) in RStudio. The boxplots represent first (lower), interquartile range (IQR), and third (upper) quartile.

### Retention time and drift time analysis

The retention time (RT) analysis was run on the timsTOF Pro2 mass spectrometer (Bruker Daltonics) with chromatographic separation through the Agilent Infinity 3 ultra-high resolution liquid chromatography attached to a Phenomenex 2.6 µm Polar C18 100 x 2.1 mm column. The synthetic samples were eluted on a gradient with flow rate of 0.5 µL/min and the mobile phase included H2O + 0.1% FA and ACN + 0.1%. Samples were eluted at 0.5 mL/min using the following gradient: 0-0.5 min, 5% B; 1.1 min, 25% B; 2.5 min, 40% B; 4.0 min, 99% B; 4.5 min, 99% B; 5.0 min, 5% B; 6.0 min, 5% B. The instrument was operated in positive ESI with capillary voltage 4500 V, end-plate offset 500 V, nebulizer pressure 2.2 bar, dry gas 10.0 L/min, and dry gas temperature 220 °C. MS data were acquired from m/z 20–1200 in PASEF mode. TIMS mobility was collected with a 1/K₀ range of 0.10–1.50 V·s/cm², ramp time of 100 ms, accumulation time of 100 ms, and a duty cycle of 100%. External mass and mobility calibration were performed using the Bruker ESI Tuning Mix (ESI-TOF CCS compendium). Extracted ion mobilograms of the [M+H]^+^ of each targeted ion were exported through the Bruker DataAnalysis 6.1 software. Raw data was converted to open-source “.mzML” format in PreoteoWizard^38^ MSConvert program (64-bit encoding, vendor peakPicking filter for MS1–MS2, zlib compression enabled) for input in the GNPS dashboard to export extracted ion chromatograms for RT plots.

### Institutional Review Board Statement

This study was supported, in part, by the analysis of standards against human samples. The sample from the HIV Neurobehavioral Research Center (HNRC) was approved by the Institutional Board Review of University of California San Diego (#172092). The sample from the osteoarthritis cohort was approved by the Institutional Board Review of University of California San Diego (#161474).

## Contributions

VD led acquisition, analysis, interpretation, and visualization of the data and writing. VCL contributed to data acquisition and interpretation. KV contributed to data acquisition. HMR contributed to data analysis and interpretation. CXW contributed to data collection and interpretation. KZ contributed resources. DS contributed resources and supervision. PCD contributed resources, supervision, and conceptualization.

## Code and data availability

Code, data tables, and other supplementary material can be found at https://github.com/victoriadeleray/3OH_N-acyl_amides. The public spectral library can be found and downloaded at https://external.gnps2.org/gnpslibrary as 3-HYDROXY-ACYL-AMIDES-LIBRARY or Zenodo^19^ at https://zenodo.org/records/17488401. Raw data from the multiplexed synthesis reactions and retention time analyses can be accessed in GNPS/MassIVE at https://massive.ucsd.edu/ProteoSAFe/dataset.jsp?task=a682622f3b464c7da6b130c62cd4c32e under accession MSV000095216.

## Acknowledgements

This work was supported, in part, by the National Institute of General Medical Sciences, R01GM107550; and the National Science Foundation [DBI-2152526]. We would also like to thank The HIV Neurobehavioral Research Center (HNRC) which is supported by award P30MH062512 from NIMH, the Guma lab supported by the Krupp Endowed Fund, and Dr. Jessica L. Metcalf, who provided samples but did not participate in the analysis or writing of this manuscript.

